# Alcama expressed in blood retina barrier and Muller glia is involved in zebrafish retina regeneration

**DOI:** 10.64898/2026.08.24.746827

**Authors:** Shiju Thomas Michael, Kristin Allan, Michael Rini, Rose DiCicco, Michael Ramos, Alex Yuan

**Author notes:** Alex Yuan, MD PhD, Cole Eye Institute, Cleveland Clinic, 2022 E 105th St, Cleveland, OH 44106, USA. These authors contributed equally (Shiju Thomas Michael, Kristin Allan).

## Abstract

Activated leukocyte cell adhesion molecule A (Alcama) plays a role in axonal guidance, cell differentiation, and retinal lamination in a developing retina and was identified as a marker for activated Muller glial cells in adult zebrafish. However, its spatiotemporal localization and its involvement in retina regeneration remains unclear. Here we induced focal photoreceptor damage in zebrafish using laser photocoagulation and examined the expression and localization of Alcama at different time points post lesion. Immunohistochemistry in wild type fish and *Tg(kdrl-EGFP)* fish showed Alcama localized to the blood retina barrier with increased expression in Muller glial end feet and radial processes in a regenerating retina. To confirm its role in retina regeneration, *alcama* expression was transiently knocked down using morpholinos in adult fish. Scanning laser ophthalmoscopy, Zpr1 immunostaining and EdU staining showed delayed retina regeneration in *alcama* knockdown fish, indicating a possible role for Alcama in zebrafish retina regeneration.

## Introduction

Retinal diseases such as age-related macular degeneration, diabetic retinopathy, proliferative vitreoretinopathy, vascular occlusion and inherited retinal dystrophies affect millions of people each year by inducing irreversible damage to retinal neurons, leading to vision loss (Zhou et al., 2023; Lundeen et al., 2023; Rein et al., 2022). Unlike mammals, zebrafish (*Danio rerio*) have an intrinsic ability to regenerate lost neurons and regain functional vision (Lahne et al., 2020). Whenever there is retinal injury in fish, the resident microglia and the infiltrating macrophages undergo dynamic transcriptional and morphological changes, inducing a pro-inflammatory response and acute inflammation (Mitchell et al., 2018). This effect is further potentiated by the cytokines released by the dying neurons. However, after a short acute inflammatory response there is onset of a regenerative response, which does not occur in mammals (Iribarne, 2021). Understanding this switch from a pro inflammatory phenotype to a regenerative phenotype and the mechanistic cross-talk between acute inflammation and regeneration are important. By identifying and exploring the functional role of molecules involved both in inflammation and regeneration in zebrafish, the mechanistic switch from a chronic inflammatory/fibrotic state in humans to a regenerative state in zebrafish can be explored.

Muller glial cells play an important role in zebrafish retinal regeneration. Whenever there is an injury to the retina, the Muller glial cells dedifferentiate and re-enter the cell cycle to generate a Muller glial cell and a proliferating progenitor cell. The progenitor cells migrate and differentiate into the required neuronal cell type, replenishing the lost cells (Goldman, 2014; Lahne et al., 2020). Using different retinal injury models, it has been found that the progenitor cells are able to replace lost neuronal cells across different retinal layers (Fausett and Goldman, 2006; Fimbel et al., 2007; Vihtelic and Hyde, 2000). Further, the end feet structures of Muller glia share boundaries with the retinal vascular endothelium and thus Muller glia are an integral part of the blood retina barrier. For instance, in zebrafish vascular endothelial growth factor Aa (Vegfaa) derived from Muller glial cells interact with its receptors (Flt1) and (Kdrl) on vascular endothelial cells to regulate Muller glial reprogramming (Mitra et al., 2022). Additionally, the integrity of the blood retina barrier plays a significant role in retina regeneration by acting as a checkpoint for the entry of immune cells and blood derived factors that drive retina regeneration (Leach et al., 2021; Nagashima et al., 2021).

Human activated leukocyte cell adhesion molecule (ALCAM), orthologue of zebrafish Alcama is a transmembrane glycoprotein that belongs to the immunoglobulin superfamily. It is known by other names such as CD166, zebrafish Neurolin, avian DM-GRASP, BEN and SC-1. A wide range of stem cells and differentiated cells express ALCAM and its orthologues, which includes mesenchymal cells, neuronal cells, leukocytes, epithelial cells and endothelial cells (Swart, 2002). Though it was initially identified as a molecule that activates leukocytes by binding to CD6 (Bowen et al., 1995), it was later found to interact with other ligands such as CD9, Galectin-8, S100B and Ezrin (Delgado et al., 2011). Because of its ability to interact with diverse binding partners, ALCAM is involved in several processes including T cell activation, axonal guidance, proliferation, migration and differentiation (Choudhry et al., 2011; Heffron and Golden, 2000; von Lersner et al., 2019). ALCAM present in brain endothelial cells is found to promote angiogenesis and regulates immune cell entry across the blood-brain barrier. For instance, ALCAM blockade prevented the lymphocyte/monocyte diapedesis across simulated pulmonary, brain and pancreatic tumor endothelium. Interestingly, ALCAM has been implicated in the transmigration of both lymphoid and myeloid leukocytes, unlike other cell adhesion molecules (Lécuyer et al., 2017; Cayrol et al., 2008).

In zebrafish, Alcama plays a significant role in neurogenesis and the development of the central nervous system, retina and peripheral motor neurons (Diekmann and Stuermer, 2009; Ott et al., 2001). Zebrafish embryos with an Alcama knockout showed significant reduction in overall eye size along with deformed retinal development. Further, *alcama* knockdown in zebrafish embryos disrupted the motor neuron axonal guidance during development (Diekmann and Stuermer, 2009). Alcama is studied predominantly in the developmental context in zebrafish, but one study reported Alcama expression during adult retina regeneration (Nagashima et al., 2013). In a light injury model, Alcama expression was observed in Muller glial cells associated with proliferating progenitor cell clusters. Further, retinal progenitors in the ciliary marginal zone which contributes to the continuous generation of retinal neurons also showed positive expression for Alcama. Therefore, Alcama was reported as a novel marker for dedifferentiated Muller glial cells (Nagashima et al., 2013). With this background, this study was designed to explore the expression and localization of Alcama at different stages of retina regeneration post-injury and to study if it plays a role in retina regeneration using a laser injury model.

## Results

### Zebrafish retina regenerates after laser injury

OCT guided laser injury was induced in zebrafish, and the presence of three lesion spots were confirmed using autofluorescence imaging by SLO (AFSLO) and OCT at 1-day post-lesion (dpl) (Supplementary figure 1A). This method of retinal injury has the advantage of delivering precise injury at the outer retina over a very short period. The injury is restricted to a small circular lesion, allowing more precise observation of the injured sites with respect to the margin zones and normal retina. The lesion size was monitored using AFSLO images taken at different timepoints post lesion to understand the extent of injury and regeneration over time. The three lesions positioned in relation to the optic nerve can be discerned from the AFSLO images. The lesion size decreased with time as seen by a significant decrease in lesion area at 7 dpl indicating wound healing and possible regeneration (Figure 1A). Cells in the outer nuclear layer, specifically the photoreceptors, were injured in the laser model. Apoptotic cells were observed using TUNEL staining at different time points in retina cryosections, indicated in cyan (Figure 1B). At 1 and 3 dpl photoreceptor death was evident in the outer nuclear layer in focal areas corresponding to the laser spot. By 7 dpl, TUNEL positive staining was seen adjacent to the outer segments as amorphous syncytial clusters, likely representing immune cells that have phagocytized apoptotic cells. Immediately after injury, we postulate that resident microglia are activated and release pro-inflammatory cytokines, which subsequently recruits monocytes and macrophages to the injured site. The number of proliferating cells across retinal layers at the site of each laser spot was measured using BrdU staining (Figure 1C) at different time points post lesion. The results indicated an increase in the proportion of BrdU positive cells in the outer nuclear layer (shown in red) and a decrease in the number of BrdU positive cells in the inner nuclear layer (shown in blue) over time (Figure 1D). The presence of proliferating progenitor cells indicates active retinal regeneration at the precise location of the laser lesions.

**Figure 1:**
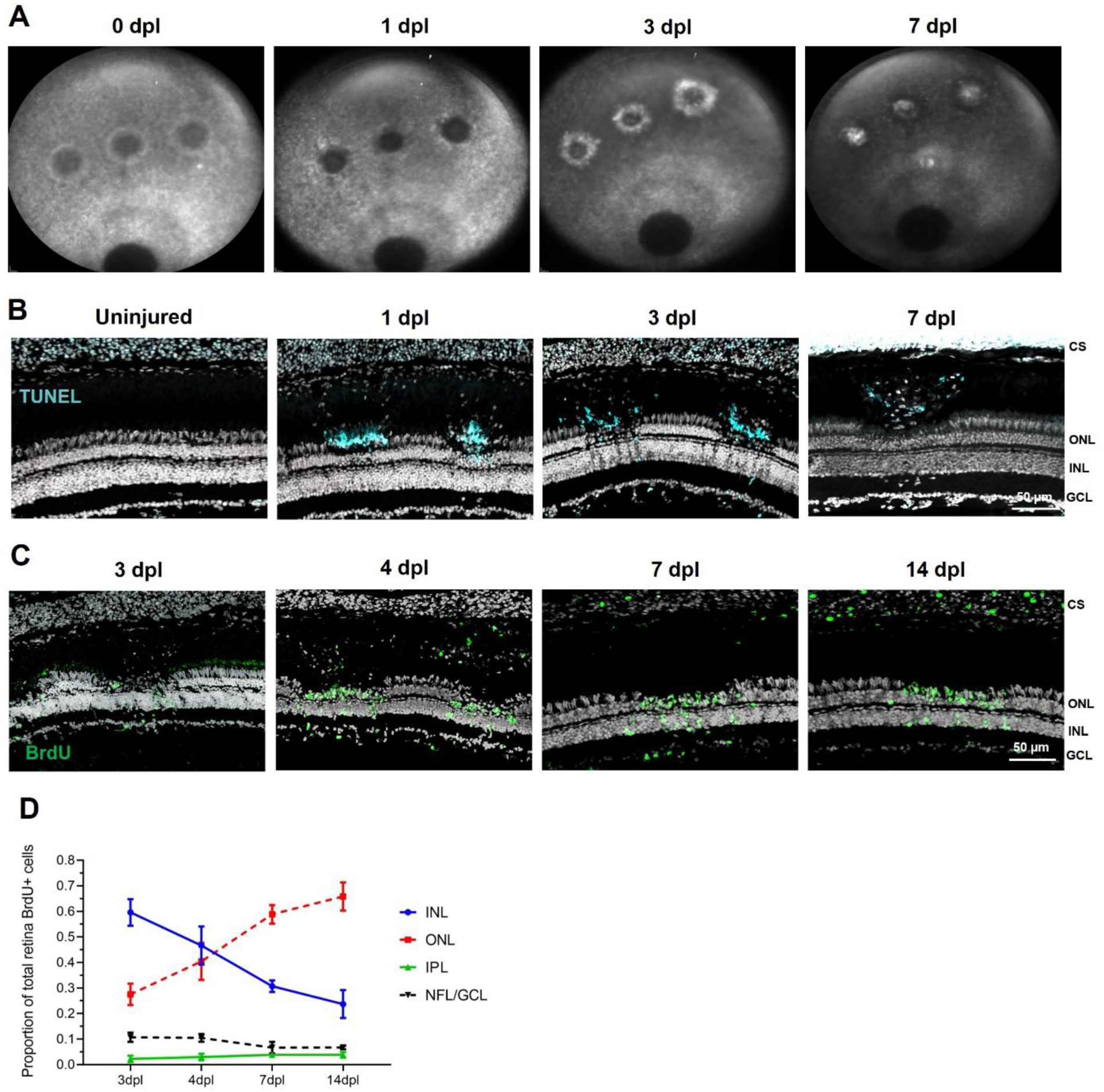
Zebrafish retina regenerates after laser injury. **(A)** Representative autofluorescence SLO images at different time points post lesion showing how the laser spots are healing and regenerating with time. **(B)** TUNEL staining (cyan, top panel) showing apoptosis in the outer nuclear layer reflecting photoreceptor loss induced by laser injury. **(C)** Representative images for retinal sections stained for BrdU (green) and counterstained with propidium iodide at different time points post lesion. **(D)** Number of BrdU-positive cells counted in each retinal layer at the center-most 3-5 sections of lesions for indicated time points and plotted in terms of percentage of total BrdU-positive cells across different retinal layers. GCL-ganglion cell layer, INL-inner nuclear layer; ONL-outer nuclear layer; IPL-inner plexiform layer; CS-choroidal stroma. Data points represent mean ± standard error of mean computed from multiple fish for each time point.

### Alcama expression in relation to other retinal cell markers

The disorganized outer nuclear layer with loss of photoreceptors at each lesion site is reflected by decreased expression of Zpr1 and Zpr3 at 1, 4, and 7 dpl. At 14 dpl, photoreceptors regenerated in and around the lesion site with positive expression for both Zpr1 and Zpr3 (Figure 2A, 2B and Supplementary figure 1B). Further, 4C4 positive microglial cells were highly expressed around the lesion in the early time points (1, 4 dpl) indicating the recruitment of activated microglia to the injured site. At 7 dpl, clusters of 4C4+ cells were observed in the outer segments near the RPE layer supporting phagocytosis and the removal of the dead neurons in the outer retina. At 14 dpl, the expression of 4C4 was similar to the uninjured retina with interspersed expression across all retinal cell layers (Figure 2A, C). Alcama (in green) showed positive expression in the choroidal endothelium, choriocapillaris, the choroid-RPE intersection (outer blood-retina barrier), and at the inner limiting membrane irrespective of the injured state. The specificity of the Alcama antibody was confirmed using Western blot and by using respective primary and secondary antibody controls in IHC (Supplementary figure 2A, B). Further, the proliferating progenitor cells that are generated from the activated Muller glial cells were identified using proliferating cell nuclear antigen (PCNA) marker, which stains the cells in the s-phase of the cell cycle. In the uninjured retina, there are few PCNA positive cells. However, at 4 and 7 dpl a greater number of cells stained positive for PCNA in the inner and outer nuclear layers at each lesion site indicating active proliferation (Figure 2A, D, and E). At 14 dpl, the number of PCNA positive cells were similar to the uninjured retina indicating the regeneration was near completion (Figure 2).

**Figure 2.**
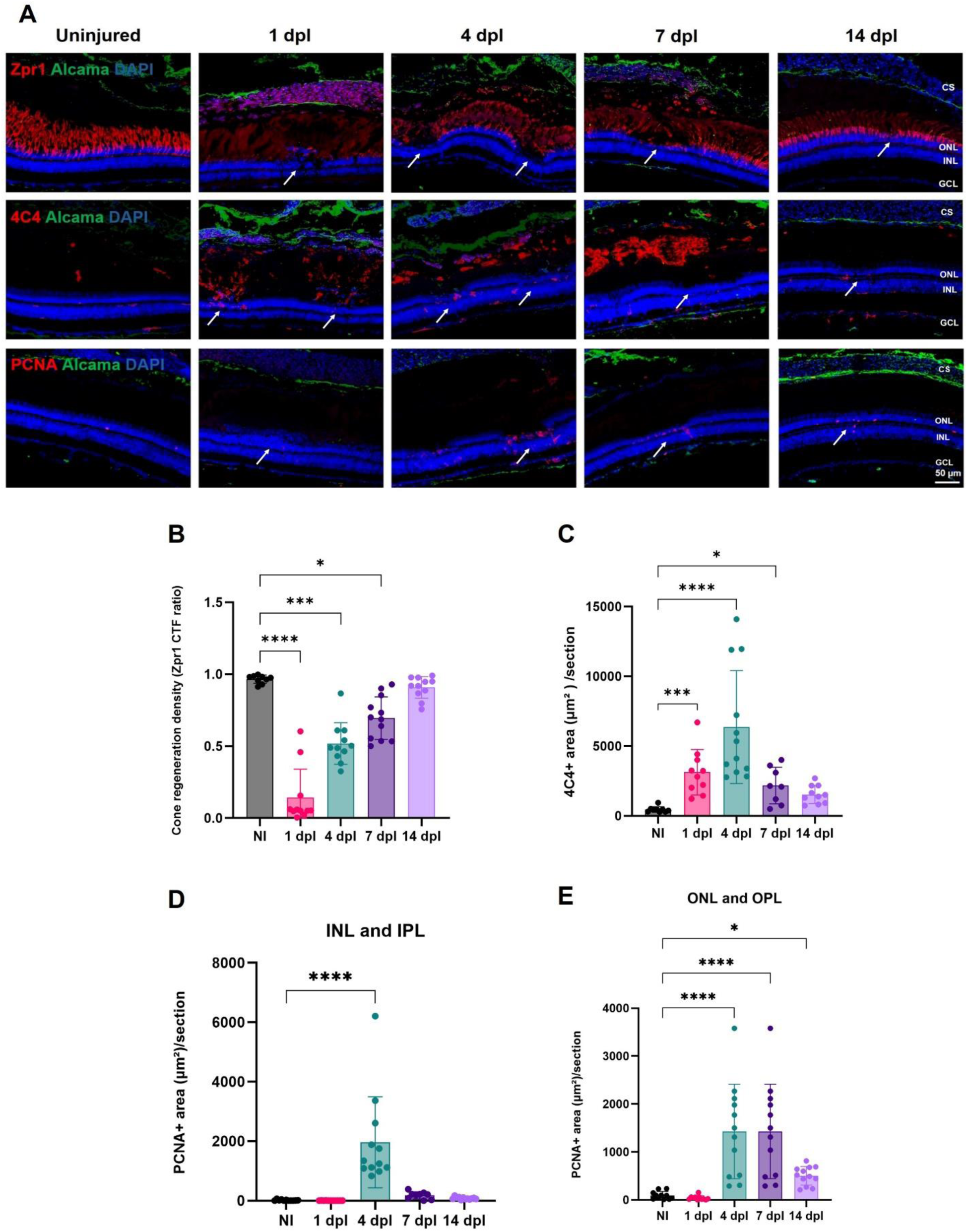
Expression of Alcama in relation to other retinal cell markers. **(A)** Representative IHC images showing Alcama (green) with Zpr1 (red) that stains the cone photoreceptors (top panel) and 4C4 (red) which stains the microglial cells and PCNA which stains proliferating cells (bottom panel). Dapi (blue) indicates the nuclear staining. Lesion spots are indicated by arrows. **(B)** Histogram showing cone regeneration density CTF ratio calculated from Zpr1 staining in fish at different time points post lesion compared to non-injured controls. Photoreceptors lost after retinal injury at earlier time points reconstituted around 14 dpl, indicating regeneration. **(C)** Histogram showing 4C4+ area per section in fish at different time points post lesion compared to non-injured controls. Activated microglia mobilize and migrate to lesion sites immediately following injury and formed giant cells, indicating phagocytosis (7 dpl). **(D, E)** Histogram showing PCNA+ area per section in fish at different time points post lesion compared to non-injured controls. Increased PCNA+ cells formed from Muller glia cells (4, 7 dpl) were responsible for regeneration. Values indicate mean ± SEM; Significance values are indicated as * p<0.05; *** p<0.001; **** p<0.0001.

### Alcama expression is increased in Muller glia during regeneration

In the uninjured retina, Zrf1 (glial fibrillary acidic protein or GFAP) positive Muller glial cells were observed in the inner nuclear layer at the inner limiting membrane and with radial extensions spanning the ganglion cell layer. Alcama expression was confined to the inner limiting membrane and nerve fiber layer, possibly in association with retinal vasculature in the uninjured retina. At 4 and 7 dpl, Zrf1 positive Muller glial cells colocalized with Alcama expression preferentially in the ganglion cell layer and in the inner plexiform layer (Figure 3A). Some of the Zrf1 positive Muller glial showed positive expression for Alcama in the radial extensions confined to the laser lesions (Figure 3B). A significant increase in the number of Alcama+ Zrf1+ radial cells at 4, 7 and 14 dpl were observed compared to the non-injured controls (Figure 3C). It is interesting to note that these Muller glial radial extensions were observed between the nerve fiber and ganglion cell layers but were not seen near the inner nuclear layer. Therefore, we believe that this increased Alcama expression is confined to activated Muller glial end feet and the distal projections of the Muller cells, primarily around the vessels in the regenerating retina.

**Figure 3.**
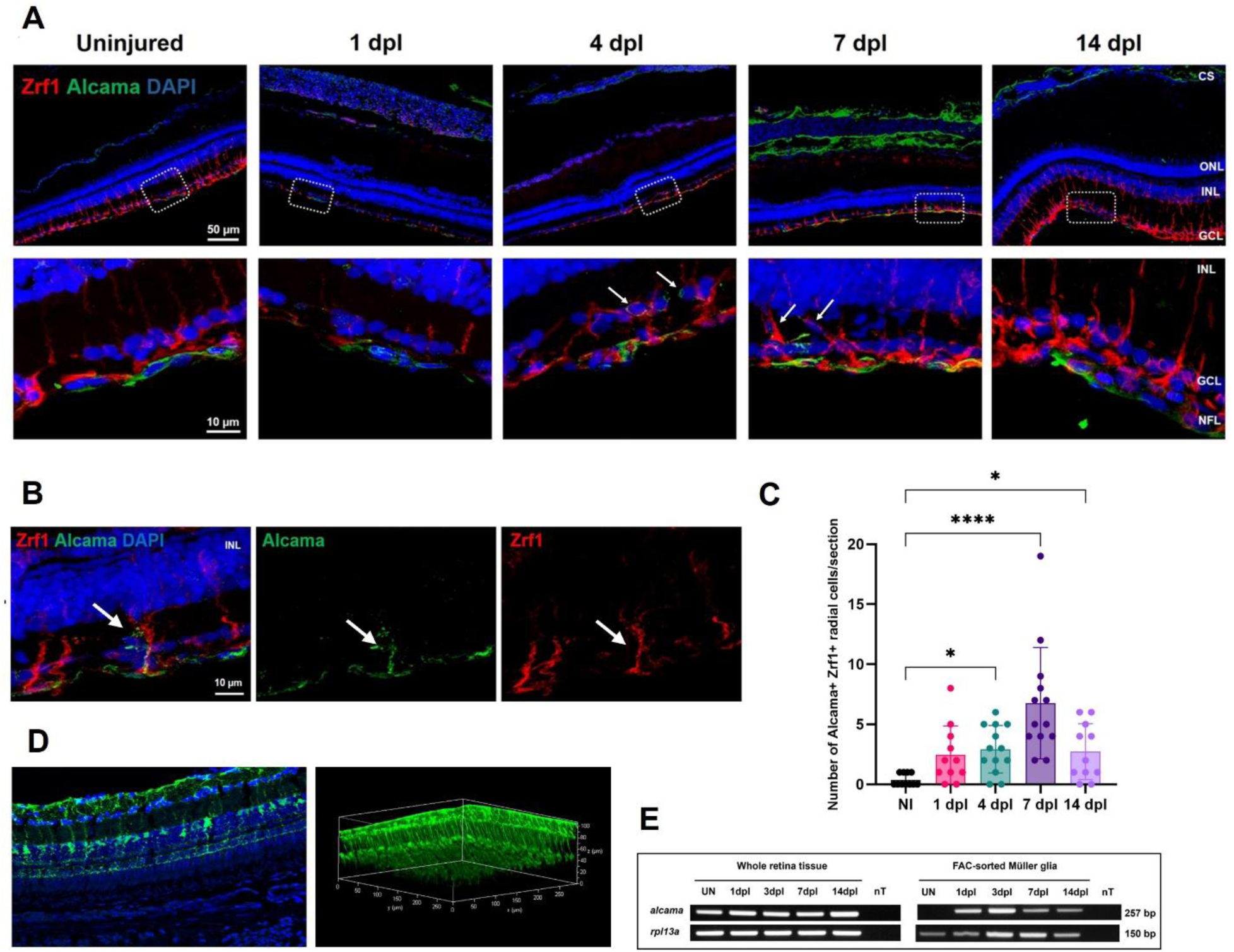
Alcama expression is increased in Muller glia during regeneration. **(A)** Representative IHC images showing Zrf1 staining of Muller glial cells (red) with Alcama (green) and Dapi (blue). At 4 and 7 dpl, Zrf1 positive Muller glial cells expressed Alcama (indicated by arrows), preferentially at the Muller glial end feet and extending radially into inner retina at 7 dpl (bottom panel). **(B)** Representative 7 dpl images showing individual channels for Alcama (green) and Zrf1 (red). Of the four Zrf1+ Muller glial cells just beneath the lesion in this representative image, one showed positive expression for Alcama. **(C)** Histogram showing number of Alcama+ Zrf1+ radial Muller glial cells per section in fish at different time points post lesion compared to non-injured controls. Values indicate mean ± SEM; Significance values are indicated as * p<0.05; **** p<0.0001. **(D)** Three-dimensional reconstruction images of flat-mounted retina from *Tg(apoe:EGFP)* zebrafish (300 x 300 x 100 μm). The GFP-positive, elongated processes spanning nearly the entire thickness of the retina exemplify Muller glia morphology, indicating the specificity of the ApoE promoter for Muller glia in the zebrafish retina. **(E)** RT-PCR analysis of whole retina and FACS-sorted Muller glial cells showing alcama expression in relation to rpl13a. nT, no template negative control. Muller glial cells were isolated via fluorescence activated cell sorting (FACS) from cell suspensions of 6-8 retinas each that were either uninjured (UN) or injured and collected at various days post lesion (dpl). Representative image of three independent experiments. Alcama expression was specifically upregulated in Muller glia in response to injury in the zebrafish retina.

To confirm the expression of Alcama in activated Muller glia, *alcama* mRNA expression was observed in FACS-sorted Muller glial cells isolated from *Tg(apoe:EGFP)* fish. Muller glia is robustly labelled with green fluorescent protein (GFP) in this line (Allan et al., 2020). In the uninjured retina, *alcama* mRNA expression was not detectable in Muller glia. After injury, the Muller glial cells showed increased *alcama* mRNA expression (Figure 3B, C).

### Alcama localized to blood retina barrier

To confirm localization of Alcama to retinal vasculature, *Tg(kdrl:EGFP)* fish was used which stains the kinase insert domain receptor like protein expressed in vascular endothelial cells. *Tg(kdrl:EGFP)* fish were laser injured and observed for Alcama expression at different time points post lesion (Figure 4). In both the uninjured and injured retina, Alcama expression co-localized with the Kdrl protein in the retinal vessels, choroidal vasculature, and choriocapillaris. This suggests that Alcama is part of the blood retina barrier (BRB) in zebrafish. In addition to the retinal vessels, Alcama expression was observed extending from the vessels along the inner limiting membrane and sometimes along radial extensions into the retina.

**Figure 4.**
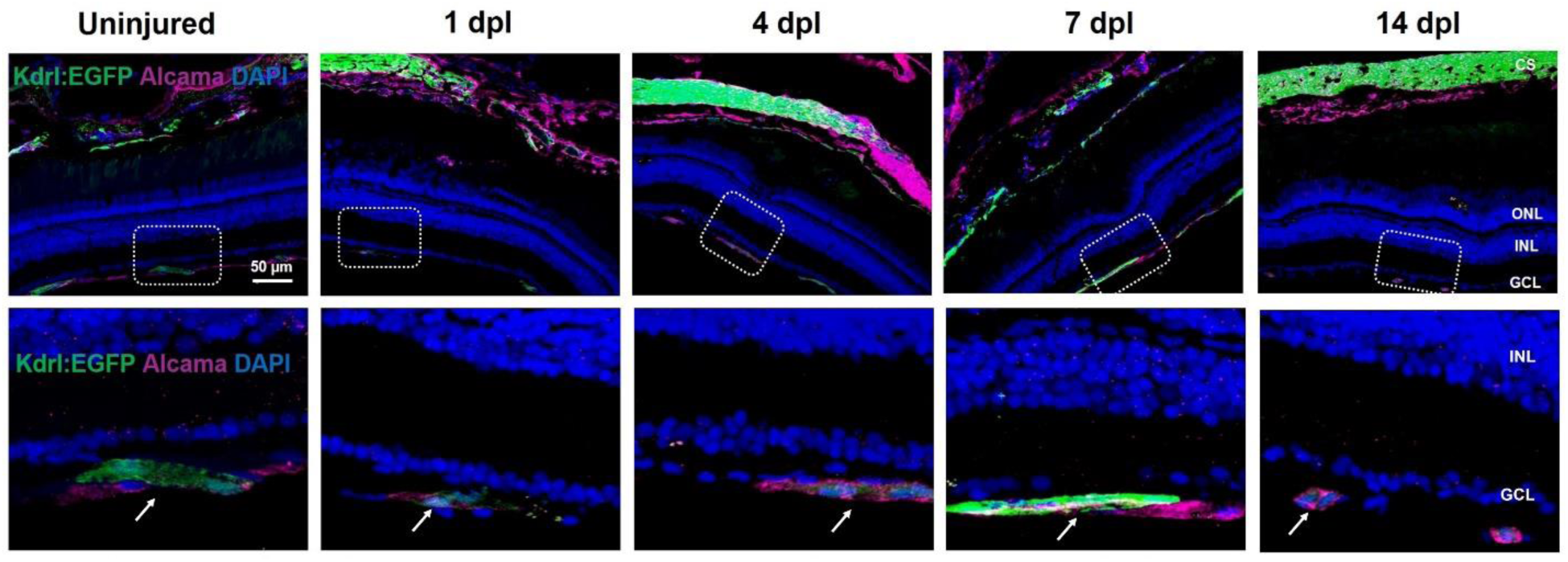
Localization of Alcama to blood retina barrier. Representative IHC images showing Alcama (magenta) with kdrl:EGFP (green) at different time points post lesion. Kdrl is a marker for vascular endothelial cells. Alcama co-localized with Kdrl in choroidal vasculature (top panel). High magnification images (bottom panel) showing colocalization of Alcama with Kdrl in the retinal vasculature in both the injured and uninjured retinas, indicated by arrows.

### Inhibition of Alcama delayed retina regeneration

Alcama is known to play a role in eye development specifically in axonal guidance and cellular patterning. Fish embryos with Alcama knockdown showed reduced eye size, deformed retina, and decreased number of cells in the INL and GCL. Therefore, this study used a transient Alcama morpholino (MO) knockdown experiment to study the role of Alcama in adult zebrafish during retinal regeneration. Morpholinos were tagged with lissamine, a positively charged fluorescent molecule, that helps in tracking the morpholinos post injection and facilitates the movement of morpholinos during electroporation for improved integration into retinal cells. Flat mounts and retina cryosections confirmed the integration of morpholinos across different cellular layers of retina (Figure 5A). The optimal concentration of Alcama MO to inhibit Alcama expression was found by titrating different Alcama MO concentrations in fish embryos with respect to the control morpholinos. Jess protein simple results showed Alcama MOs at a threshold of 2.5 ng were able to knockdown Alcama protein expression (Figure 5B and C).

**Figure 5:**
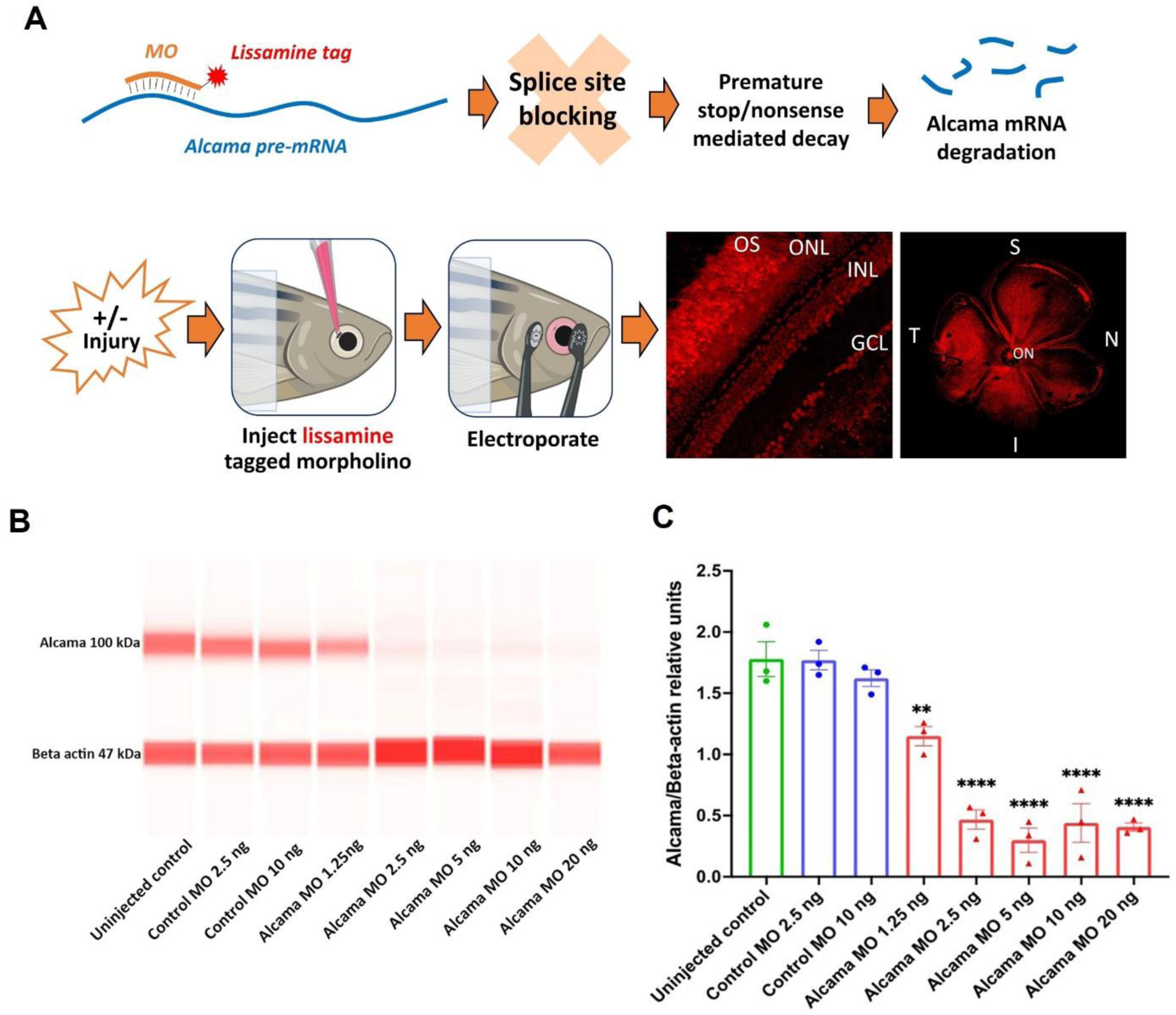
Alcama knockdown using lissamine tagged morpholinos. **(A)** Flat mount and histological retina sections showing lissamine tagged morpholino (Red) delivered to all retinal layers. **(B, C)** Jess protein simple showing Alcama expression in fish embryos treated with morpholinos at different concentrations. Values indicate mean ± SEM; Significance values are indicated as ** p<0.01; *** p<0.001 **** p<0.0001.

Using AFSLO imaging, we took longitudinal measurements of each laser lesion in animals treated with *alcama* MO and control “scrambled” MO. AFSLO images were taken every alternate day following laser injury up to 14 days (Figure 6A), and the lesion area was measured using Image J. The percentage lesion area relative to its baseline measurement was computed to understand how the lesion size changed over time (Figure 6B). The lesion area decreased with time in both Alcama MO and control MO treated groups indicating regeneration in both groups. However, there was a significant difference in the lesion area at 8, 10, and 12 dpl in Alcama MO treated samples compared to the control MO. This indicated that Alcama knockdown attenuated the overall retinal regeneration rate.

**Figure 6:**
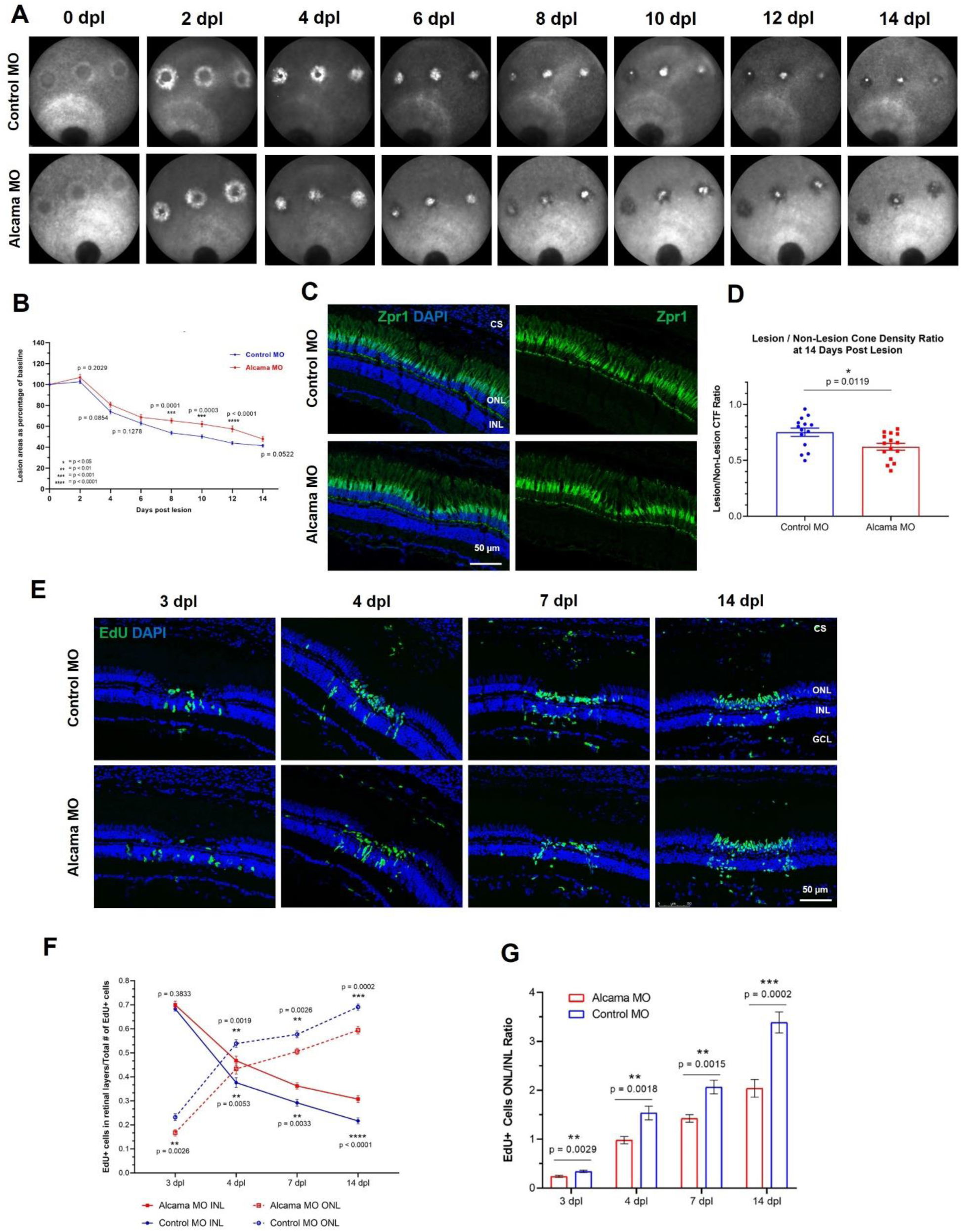
Inhibition of Alcama delayed retina regeneration. **(A)** Representative autofluorescence SLO images showing lesions in fish treated with *alcama* morpholino compared to control morpholino treated fish. **(B)** The lesion area was quantified and represented as mean ± SEM. Significance values are indicated as * p<0.05; ** p<0.01; *** p<0.001; **** p<0.0001. **(C, D)** Representative IHC images stained for Zpr1 (Green) and DAPI (Blue) at 14 days post-lesion. Cone regeneration density is represented as lesion/non-lesion ratio. Values represent mean ± SEM. Significance values are indicated as * p<0.05. **(E)** Representative images from EdU lineage tracing assay. Each image shows a single z-stack of a single 10 μm section. MO - morpholino. **(F)** Total EdU-positive cells quantified within all retinal layers, plotted in terms of number per section. n = 14-18 lesions at each time point in both treatment groups, with 6 fish represented in all groups. INL - inner nuclear layer; ONL - outer nuclear layer. **(G)** ONL/INL ratios of EdU-positive cells over time after injury. Error bars represent standard error of the mean. Adjusted p-values obtained from multiple t-tests comparing the control and Alcama morpholino treatments with Holm-Sidak correction. Significance values are indicated as * p<0.05; ** p<0.01; *** p<0.001 **** p<0.0001.

To understand the effect of Alcama knockdown on cone regeneration, Zpr1 staining was performed in fish treated with either Alcama MO or control MOs. Cone regeneration density was quantified by measuring the Zpr1 signal intensity in both the lesion and non-lesion areas and represented as a ratio. At 14 dpl, there was a significant difference in cone regeneration density between Alcama MO and control MO groups, indicated that Alcama knockdown resulted in decreased cone regeneration (Figure 6C and D). To further define the effect of Alcama knockdown on retinal regeneration, EdU lineage tracing was performed and the number of EdU+ proliferating cells (in green) in different retinal layers at different time points was quantified (Figure 6E and F). EdU labelling was performed at 3 dpl, followed by imaging at 3, 4, 7 and 14 dpl. The number of EdU+ cells in the outer nuclear layer in the Alcama MO treated group was significantly lower compared to the control MO group at 3 dpl (p=0.0026), 4 dpl (p=0.0019), 7 dpl (p= 0.0026) and 14 dpl (p= 0.0002). In contrast, EdU+ cells were increased in the inner nuclear layer in Alcama MO treated group compared to the control MO treated group at 4 dpl (p=0.0053), 7 dpl (p= 0.0033) and 14 dpl (p< 0.0001) (Figure 6F). This suggested a greater population of proliferating cells remained in the INL following Alcama knockdown. The ONL/INL ratio of EdU+ cells indicated a significant decrease in EdU+ cells at the ONL relative to the INL at all time points in Alcama MO treated eyes compared to control MO, which supports Alcama’s role in regeneration of the injured photoreceptors in our laser model.

## Discussion

This study reports for the first time that Activated leukocyte cell adhesion molecule A (Alcama) is associated with the zebrafish blood retinal barrier (BRB) and is involved in retina regeneration. We found Alcama protein expression in zebrafish retinal vasculature, choroid, and choriocapillaris using IHC and report that it may be an integral part of inner and outer BRB. Alcama is consistently expressed in retinal and choroidal vessels in both the injured and uninjured retina. Following injury, Alcama expression is seen along radial extensions which stain positive for Zrf1, a Muller glia marker, confirming the previous findings from Dr. Pamela Raymond’s group. Additionally, this study finds that in the retina, Alcama expression is confined predominantly to Muller glial end feet and is proposed to be a part of the inner limiting membrane. Alcama knockdown experiments clearly showed a delay in regeneration compared to controls, suggesting a role for Alcama in zebrafish retinal regeneration.

Immunohistochemistry observations of wild-type zebrafish with/without laser injury showed expression of Alcama in retinal vessels, choroidal vessels, and choriocapillaris. The injury state and regeneration did not appear to alter the intensity of Alcama expression at the vascular endothelium, but the pattern of expression in Muller glia seemed to extend beyond the inner limiting membrane to radial extensions. *Tg(kdrl:EGFP)* fish which stains an ortholog to vascular endothelial growth factor receptor 2 in retinal and choroidal vessels further confirmed the co-localization of Alcama to the vascular endothelium (Figure 4). Thus, we propose that Alcama may be an integral component of the BRB with possible functions in mediating the transmigration of leukocytes and maintaining BRB integrity. To our knowledge, no previous studies have reported the presence of Alcama at the BRB. However, the human/mouse orthologue of Alcama (ALCAM) was previously found to be an important component of the blood brain barrier (BBB). ALCAM was localized to cholesterol enriched membrane microdomains called lipid rafts on the surface of brain endothelial cells (Mañes et al., 2006). ALCAM expression in brain endothelial cells was found to be an important mediator of leukocyte transmigration into the central nervous system (Cayrol et al., 2008). For instance, in an *in vitro* model of the BBB, ALCAM expressed in human endothelial cells was involved in transmigration of human CD4^+^, CD19^+^, and CD14^+^ leukocytes. Confirming further, ALCAM blockade in a mouse model of multiple sclerosis impacted the migration of CD3^+^, CD4^+^ and Mac-1^+^ leukocytes into the CNS, decreased the area of demyelination, reduced the number of inflammatory lesions, and eventually improved the clinical severity of the disease (Cayrol et al., 2008). The involvement of ALCAM in immune cell transmigration is also observed in other tissues. In rat pulmonary endothelial cells, ALCAM was found to mediate the migration of THP-1 monocytes (Masedunskas et al., 2006) and regulate T-cell entry in pancreatic tumors (Nummer et al., 2007). Further, in simulated pulmonary and pancreatic endothelium, ALCAM is involved in transmigration of leukocytes (Cayrol et al., 2008; Masedunskas et al., 2006; Nummer et al., 2007). Therefore, we speculate a similar role for Alcama in the BRB by mediating the transmigration of leukocytes and facilitating the acute inflammatory phase needed for successful regeneration.

Alcama present in the retinal vascular endothelium may be providing barrier integrity to the BRB. In higher mammals, the loss of ALCAM was found to compromise the permeability of BBB. For example, mice with ALCAM KO showed an increased permeability of the BBB and developed severe experimental autoimmune encephalomyelitis lesions (Lécuyer et al., 2017). ALCAM indirectly links the junction molecules to the actin cytoskeleton and thus loss of ALCAM affects the integrity of the BBB. In primary cultures of brain microvascular endothelial cells, ALCAM is found to have a direct association with intracellular TJ adapter molecules, such as ZO-2 and cingulin (Lécuyer et al., 2017). At the intracellular level, ALCAM is linked to the actin cytoskeleton and is involved in cell adhesion, partly mediated by actin polymerization (Tudor et al., 2014). In addition to cell adhesion, ALCAM is also involved in various cell migration processes such as neurite outgrowth, neurogenesis, blastocyst implantation and melanoma invasion (Swart, 2002).

Muller glial cells specifically upregulated the expression of Alcama post-injury at specific time points during regeneration. This confirmed the only previous finding showing Alcama expression localized to activated Muller glial cells (Nagashima et al., 2013). Our results are also in agreement with a recent single cell RNA sequencing study reporting upregulation of Alcama in zebrafish Muller glia after outer nuclear layer injury although not in mice and humans (Hoang et al., 2020). Additionally, our IHC observations confirmed that Alcama expression was predominantly confined to the Muller glial end feet and may be a part of the inner limiting membrane. Mammalian ALCAM is found to co-localize with VE-cadherins and N-cadherins at lateral membrane domains of endothelial cell junctions (Ofori-Acquah et al., 2008). N-cadherin^+^ retinal progenitor cells were found to form regeneration niche around N-cadherin^+^, Alcama^+^ Muller glial cells in zebrafish (Nagashima et al., 2013). Therefore, Alcama expressed in Muller glial cells may be involved in cell migration.

Lesion area measurements from AFSLO images and Zpr1 IHC showed a delay in regeneration in fish with Alcama MO knockdown compared to control MO. Furthermore, zebrafish with Alcama MO knockdown showed a significant difference in the number of EdU^+^ proliferating cells in both the inner and outer nuclear layers at multiple time points (4, 7, 14 dpl) compared to control MO (Figure 6E, F, G). Overall, the findings from Alcama knockdown experiments suggest that Alcama plays a novel role in retinal regeneration. Prior studies on zebrafish Alcama are primarily in the developmental context showing its role in axonal guidance and cellular patterning. Fish embryos with Alcama knockdown were found to have reduced eye size, deformed retina with decreased number of cells in the INL and GCL (Diekmann et al., 2009). Based on the previous literature and our current findings, we propose that Alcama expressed in BRB and Muller glial end feet mediate the transmigration of leukocytes into the retina during injury. Thus, it helps in maintaining the acute inflammatory phase following injury and may also play a role in retinal regeneration.

## Materials And Methods

### Zebrafish maintenance

Adult zebrafish were housed in 3L and 10L tanks on an Aquatic Habitats recirculating water system (Pentair; Apopka, FL) in a 14:10 hr light:dark cycle at 28.5°C and fed with brine shrimp and commercial flake food. All experiments and procedures involving animals were approved by the Institutional Animal Care and Use Committee at the Cleveland Clinic and in accordance with the guidelines by the ARVO statement for the use of animals in ophthalmic and vision research. Transgenic fish lines - *Tg(kdrl:EGFP)* was kindly donated by Dr. Matsuoka. For enucleation, fish 6-18 months old were dark adapted overnight and euthanized by placing one at a time in a divided tank with ice on the opposite side, until unresponsive to tail pinch. For procedures requiring anesthesia, fish were placed in 0.16 mg/mL tricaine in system (fish) water until unresponsive to external stimuli.

### Retinal injury by laser photocoagulation

Retinal injury/lesion was induced using a laser photocoagulation method as described previously (DiCicco et al., 2014). Briefly, the dark-adapted fish were anesthetized using tricaine and a polymethylacrylate (PMMA) contact lens (Ø = 5.2 mm, r = 2.70 mm, center thickness = 0.4 mm; Cantor & Nissell, Ltd., Northamptonshire, UK), was placed on the left eye. Laser photocoagulation was carried out using an OCT imaging system with 7 μm axial resolution (SDOIS 840HR; Bioptigen, Durham, NC, USA) fitted with a 532 nm diode laser (Oculight GL; Iridex, Mountain View, CA) using wide-field objectives (≥50° field of view). The laser was incorporated into the OCT imaging path with a custom beam combining module, which consists of a dichroic beamsplitter (685-nm long-pass; Semrock, Inc., Rochester, NY), relay optics (Thorlabs, Newton, NJ), and a modified 200 μm multimode optical fiber. An adjustable aperture set to 7.3 mm was used to deliver a 300 ms pulse of 42-47 mW output power for each lesion. Laser spots were targeted with the en face image and rotating the fish accordingly on a 5-axis positioning stage (Model AIM-RAS; Bioptigen, Inc.). To simplify histological analysis and identification of lesions, three laser lesions were placed superior to the optic nerve. When retinas were collected for expression analysis, as many lesions as allowed by the physical and optical restraints of the system and time of anesthesia were placed superior to the optic nerve, typically around 45 lesions.

### BrdU treatment and staining

Three days post-lesion (dpl), fish received two intraperitoneal injections of Bromodeoxyuridine (BrdU) (0.25 μmol/g body weight), two hours apart. Eyes were collected and retinal sections were stained for BrdU - 4 hours after the second injection for the 3-day time point. Samples were collected and processed for subsequent time points (4, 7 and 14 dpl) as well. Enucleated eyes were placed in optimal cutting temperature media, flash frozen in a 2-methylbutane bath over liquid nitrogen and stored at-80°C before cryo-sectioning into 10 μm sections using CFSA 0.5x slides (Cat #39475285; Leica Microsystems Inc, IL, USA) and adhesive tape (Cat #39475214; Leica Microsystems Inc, IL, USA). Retina sections were stained using a BrdU Labeling and Detection Kit following the manufacturer’s protocol (Cat #10280879001; MilliporeSigma, MD, USA) and counterstained with propidium iodide.

### Immunohistochemistry and confocal microscopy

Zebrafish eyes were enucleated immediately post-euthanasia and fixed in 4% paraformaldehyde overnight at 4°C. To cryoprotect the eyes, each sample was treated with a gradient sucrose of 10%, 20% and 30% in 1x PBS for 2 hours each at 4°C with gentle mixing on a rotor. Samples were then incubated overnight in 1:1 solution of 30% sucrose and OCT at 4°C. Tissues were then embedded in OCT and stored at-80°C until cryo-sectioning. 10 μm sections were prepared using a cryotome (Leica CM1950) and mounted on superfrost slides (Leica, Cat #1255015) and stored at-20°C. On the day of staining, slides were allowed to thaw for 15 mins at room temperature and rehydrated by treating with 1X PBS twice for 5 mins each. Autofluorescence was quenched by incubating the slides in a bleaching buffer (1x SSC, 5% peroxide and 5% Formamide) for 10 mins. Slides were then incubated with the blocking buffer (5% Goat/Donkey serum and 5% BSA in PBS) for 1 hour at room temperature. Primary antibodies at different dilutions were added to the slides and incubated overnight at 4°C. The details about the primary antibodies and the dilutions used in this study can be found in Table 1. The following day, slides were washed thrice with PBSTD (1xPBS with 0.1% Tween20 and 0.1% DMSO), 5 mins each wash. After washing, slides were incubated with corresponding secondary antibodies for 1 hour at room temperature, the details of which can be found in Table 2. The slides were then washed with PBSTD thrice and incubated with DAPI (Cat# 62248; ThermoFisher Scientific, MA, USA) for nuclear staining. Finally, slides were mounted using few drops of Fluoromount-G (Cat# 0100-01; SouthernBiotech, AL, USA) and stored at 4°C until imaging. Immunofluorescence images were captured using the Leica TCS SP8 confocal scanning microscope (Leica Microsystems, Europe). Three-dimensional z-stack images were obtained by scanning the entire thickness of retinal sections and processed with the 3D Viewer in the Leica Application Suite X Software.

### RNA extraction and RT-PCR

For whole retina analysis, retinas were dissected out and transferred to a nuclease-free Eppendorf tube held on dry ice, and stored at-80°C. To extract RNA, cold TRIzol reagent (Invitrogen) was added before homogenizing with handheld motorized pestle, and the manufacturer’s protocol was followed to obtain RNA with A260/280 around 1.8 and A260/230 around 2. For FAC-sorted Muller glia, *Tg(apoe:EGFP)* retinas were dissected, dissociated, and prepared for flow cytometric cell sorting, as described previously (Allan et al., 2020). Muller glial cells were sorted directly into Buffer RLT at 4°C, and RNA was extracted using the RNeasy Micro Kit (Cat #74004; Qiagen) following the manufacturer’s protocol, with final RIN values of at least 7. Reverse transcription was carried out with oligo(dT) primers, random hexamers, and SuperScript IV Reverse Transcriptase (Cat #18090010; ThermoFisher Scientific, MA, USA) to generate cDNA following the manufacturer’s protocol. Subsequent PCR reactions (RT-PCR) were performed with EmeraldAmp GT PCR Master Mix (Cat #RR310A; Takara Bio Inc.) and gene-specific primers (alcama F: 5’-AGGCACAGAAAGATGATCCG-3’, alcama R: 5’-ACATTCCCCCAACTGCGTC-3’; rpl13a F: 5’-TCTGGAGGACTGTAAGAGGTATGC-3’; rpl13a R: 5’-AGACGCACAATCTTGAGAGCAG-3’).

### Morpholino treatments

Custom-designed lissamine-tagged morpholinos (Gene Tools LLC, OR, USA) were received as a lyophilized powder and reconstituted in sterile, deionized water to a 3 mM working solution. The anti-*alcama* morpholino (5’-TGTGTTTAAGCTATGCTTACTGTGA-3’) was designed to target and bind a splice site acceptor sequence between exon 5 and intron 5, causing introduction of a premature stop codon. Scrambled oligos (5’-CCTCTTACCTCAGTTACAATTTATA-3’) with no known binding site in the zebrafish genome/transcriptome was used as morpholino control. Morpholinos were delivered intravitreally using a previously described protocol with slight modifications (Thummel et al., 2011). Briefly, anesthetized fish was placed on paper towels soaked in system water under a dissecting microscope at 4-5x magnification. The outer cornea was first removed using one pair of Dumont #1 forceps (or similar), one to stabilize the eye and another to pull off the outer cornea. The tip of a 15-degree angle stab blade (LaserEdge Plus, PL7515) was used to make an incision in the superior-temporal region of the Iris. A pulled 1 mm capillary needle was then used to draw up 0.5 μL of 3 mM morpholino solution by capillary action. The needle was inserted into the incision and injected intravitreally using a microinjector with an injection time of 0.9 ms (Nikon PLI-188, Garden City, NY). The eye was gently rotated within the socket with platinum-plated electroporation tweezers (CUY650P3), with the negative electrode oriented behind the eye. A 75 V pulse was then delivered for 50 ms duration (BTX ECM 830 Square Wave Electroporation System, Harvard Apparatus; connection cables for electroporation tweezers: C115SCB-2).

### Analysis of *alcama* morpholino efficiency in zebrafish embryos

To check the efficiency of the *alcama* MO, different concentrations were prepared by diluting in nuclease free water. Morpholinos at different concentrations were administered into the center of the yolk at single/2-cell stage, to reduce the chance of secondary effects. Alcama MOs were used at different quantities ranging from 1.25 ng, 2.5 ng, 5 ng, 10 ng and 20 ng, whereas the scrambled control MOs were used at 2.5 and 5 ng. A minimum of 40-50 embryos were injected per morpholino concentration. The MO injected embryos were cultured in embryo water in an incubator at 29°C. Uninjected zebrafish embryos at 4 days post fertilization were used as controls.

### Jess protein simple to quantify Alcama protein expression

Jess automated western nanoassay system (ProteinSimple, Bio-Techne, CA, USA) was used with internal standards to quantify Alcama protein expression in zebrafish embryos treated with different concentrations of Alcama morpholinos. Protein lysates were prepared using Pierce™ RIPA lysis buffer (Cat. #89900, ThermoFisher Scientific, MA, USA) and the extracted proteins were quantified using Pierce™ BCA protein assay kit (Cat. #23227, ThermoFisher Scientific, MA, USA). The protein lysates were diluted with 0.1x sample buffer and mixed with fluorescent standards and 400 mM dithiothreitol to reach a final concentration of 0.6 μg/μl. The protein samples were mixed and denatured at 95°C for 5 mins and stored on ice. The 12-230 kDa fluorescence separation module was used in this study with the 25-well capillary system. The assay plate was set up as recommended in the instrument manual. Antibody diluent, Alexa fluor conjugated anti-rabbit and anti-mouse secondary antibodies and luminol peroxide mix were used from ProteinSimple. Primary antibodies corresponding to Zebrafish Alcama (Cat. # GTX128399, GeneTex, CA, USA) and beta-actin (Cat #sc-47778, Santa Cruz Biotechnology, TX, USA) were used at 1:10 and 1:25 dilutions respectively. The samples were separated at 375 V for 25 min and subjected to blocking with the antibody diluent for 30 mins, 90 min of primary antibody incubation and 60 min of secondary antibody incubation. The data obtained from Jess were virtual blots with bands representing the protein of interest and the electropherograms representing the signal intensity of each band. The area corresponding to each band was also computed using the Compass simple western software.

### Confocal scanning laser ophthalmoscopy (cSLO) imaging and quantification

Polymethylacrylate (PMMA) contact lens (Ø = 5.2 mm, r = 2.70 mm, center thickness = 0.4 mm; Cantor & Nissell, Ltd., Northamptonshire, UK), was placed over the anesthetized fish and cSLO images were obtained with a HRA2 system (Heidelberg Engineering, Carlsbad, CA) equipped with wide-field objective (≥50° field of view). Lesion areas were quantified with Image J software. Lesions were outlined with the freehand tool and area quantities obtained using the measurement function. The line drawing tool was used to draw a straight line between the centers of the temporal lesion (TL) and nasal lesion (NL), aligning the ends of the line with the labels of each lesion, which automatically appear at the centroid of the drawn lesion areas. Images were standardized using the TL-NL distance measurement to minimize differences in transverse magnification. This linear measurement was squared to translate a linear measurement to area measurement. All time points were normalized to day 0 to obtain a scaling factor. This scaling factor was multiplied by the raw area measurement to get the final scaled lesion area measurements used for statistical analysis. The average measurements from three independent graders, masked to the treatment arms, were used for final analysis (Supplementary figure 2C).

### Zpr1 immunofluorescence quantification

Images of Zpr1-stained sections were used to quantify relative cone density using Image J software. Integrated density (ID) and area (A) measurements were obtained for both lesion and non-lesion zones; average background fluorescence (ABF) measurement was made by computing the background mean fluorescence measurements obtained above and below the stained region of interest. As a readout for cone density within and outside a lesion, corrected total fluorescence (CTF) was calculated for both zones: lesion CTF = ID(lesion) – [A(lesion) × ABF]; non-lesion CTF = ID(non-lesion) – [A(non-lesion) × ABF]. The CTF ratio was then calculated for an individual image: CTF ratio = Lesion CTF ÷ Non-Lesion CTF. These measurements were averaged for sections spanning an entire lesion, about 10-12 sections per lesion, to obtain an individual data point indicating the density of regenerated cones relative to the baseline cone density in the region of interest (Supplementary figure 2D).

### EdU staining and quantification

Three days post lesion, 20 μL of 20 mM EdU solution was injected intraperitoneally with a 33g needle. Eyes were collected four hours later for a 3-day time point, or at subsequent time points beyond three days. Cryosections were obtained as previously described and stained for EdU using the Click-It EdU Imaging Kit as recommended by the manufacturer’s protocol (Invitrogen, C10337). Samples were washed in 1x PBS and counterstained with DAPI (Cat# 62248; ThermoFisher Scientific, MA, USA) at 1 μg/mL in 1x PBS. EdU-positive cells were counted manually from images of retinal section z-stacks (10 μm) for each of five retinal layer groupings: NFL/GCL, IPL, INL, ONL/OPL, OSZ/RPE. NFL – Nerve fiber layer; GCL – Ganglion cell layer; IPL – Inner plexiform layer; INL – Inner nuclear layer; ONL – Outer nuclear layer; OPL – Outer plexifirm layer; OSZ – Outer segment zone; RPE – Retinal pigment epithelium. If a cell spanned two layers, it was counted in the layer which contained most of the cell. This was repeated for every section encompassing an entire lesion. For each layer, EdU-positive cell counts were averaged across all sections (mean of 16 sections/lesion) to obtain one ‘n’ value, representative of a single lesion. ONL/INL ratios were obtained by dividing the average ONL cell count obtained for each lesion by the average INL cell count for the same lesion; this was repeated for each lesion as a single ‘n’ value. Total EdU-positive cell counts were obtained by dividing all EdU-positive cells counted in all layers of a lesion by the total number of sections (i.e. images) quantified for that lesion to normalize the counts.

## Statistical analysis

GraphPad Prism 11.0.1 was used for statistical analyses and preparing figures. Independent t-tests (two-tailed) were performed, with Holm-Sidak correction when multiple time points were analyzed. For IHC quantification data, one-way ANOVA with Kruskal-Wallis multiple comparison test was performed to compare different time points post lesion with the non-injured groups. Values were considered statistically significant when p value was less than 0.05. Significance values were indicated as * p<0.05; ** p<0.01; *** p<0.001 **** p<0.0001.

## Data Availability

The authors declare that the data supporting the findings of this study are available within the paper and its supplementary information. The microscopic images obtained from individual fish and the numerical data used for analysis is available upon request. Any additional information required to reanalyze the data reported in this paper are available upon request from the corresponding author.

## Acknowledgements

The authors thank the staff of Biological Research Unit in Cleveland Clinic Research for their excellent animal care. The authors are grateful to Dr. Ryoto Matsuoko for the *Tg(kdrl-EGFP)* fish line. This study was supported by the National Institutes of Health/National Eye Institute - 5T32EY024236-04, NEI1K08EY023608, and P30EY025585, and Research to Prevent Blindness Centre Award unrestricted grant to Cleveland Clinic Lerner College of Medicine.

## Author Contribution

STM: Study design, experimental procedure, data analysis, manuscript writing, manuscript editing; KA: study design, experimental procedure, data analysis; MR: experimental procedure, data analysis; RD: experimental procedure; data analysis MR: experimental procedure, data analysis; AY: Conceptualisation, study design, manuscript review, manuscript editing and funding acquisition.

Shiju Thomas Michael (STM) and Kristin Allan (KA) contributed equally for this study. Correspondence to Alex Yuan

## Supplementary information

Supplementary figures and tables are included as a separate file.

## References

Allan, K., DiCicco, R., Ramos, M., Asosingh, K. & Yuan, A. Preparing a single cell suspension from zebrafish retinal tissue for flow cytometric cell sorting of Muller glia. Cytometry A 97, 638–646 (2020).

Bowen, M. A., et al. Cloning, mapping, and characterization of activated leukocyte-cell adhesion molecule (ALCAM), a CD6 ligand. J. Exp. Med. 181, 2213–2220 (1995).

Cayrol, R., et al. Activated leukocyte cell adhesion molecule promotes leukocyte trafficking into the central nervous system. Nat. Immunol. 9, 137–145 (2008).

Choudhry, P., Joshi, D., Funke, B. & Trede, N. Alcama mediates Edn1 signaling during zebrafish cartilage morphogenesis. Dev. Biol. 349, 483–493 (2011).

Delgado, V. M. et al. Modulation of endothelial cell migration and angiogenesis: a novel function for the “tandem-repeat” lectin galectin-8. FASEB J. 25, 242–254 (2011)

DiCicco, R. M. et al. Retinal regeneration following OCT-guided laser injury in zebrafish. Invest. Ophthalmol. Vis. Sci. 55, 6281–6288. (2014).

Diekmann, H. & Stuermer, C. A. O. Zebrafish neurolin-a and-b, orthologs of ALCAM, are involved in retinal ganglion cell differentiation and retinal axon pathfinding. J. Comp. Neurol. 513, 38–50 (2009).

Fausett, B. V. & Goldman, D. A role for alpha1 tubulin-expressing Muller glia in regeneration of the injured zebrafish retina. J. Neurosci. 26, 6303–6313 (2006).

Fimbel, S. M., Montgomery, J. E., Burket, C. T. & Hyde, D. R. Regeneration of inner retinal neurons after intravitreal injection of ouabain in zebrafish. J. Neurosci. 27, 1712–1724 (2007).

Fleckenstein, M., et al. Age-related macular degeneration. Nat. Rev. Dis. Primers. 7, 31 (2021).

Goldman, D. Muller glial cell reprogramming and retina regeneration. Nat. Rev. Neurosci. 15, 431–442 (2014).

Heffron, D. S. & Golden, J. A. DM-GRASP is necessary for nonradial cell migration during chick diencephalic development. J. Neurosci. 20, 2287–2294 (2000).

Hoang, T., et al. Gene regulatory networks controlling vertebrate retinal regeneration. Science 370, eabb8598 (2020).

Iribarne, M. Inflammation induces zebrafish regeneration. Neural. Regen. Res. 16, 1693–1701 (2021).

Lahne, M., Nagashima, M., Hyde, D. R. & Hitchcock, P. F. Reprogramming Muller glia to regenerate retinal neurons. Annu. Rev. Vis. Sci. 6, 171–193 (2020).

Leach, L. L., Hanovice, N. J., George, S. M., Gabriel, A. E. & Gross, J. M. The immune response is a critical regulator of zebrafish retinal pigment epithelium regeneration. Proc. Natl. Acad. Sci. U. S. A. 118, e2017198118 (2021).

Lécuyer, M. A., et al. Dual role of ALCAM in neuroinflammation and blood-brain barrier homeostasis. Proc. Natl. Acad. Sci. U. S. A. 114, E524–E533 (2017).

Lundeen, E. A., et al. Prevalence of Diabetic Retinopathy in the US in 2021. JAMA Ophthalmol. 141, 747–754 (2023).

Mañes, S. & Viola, A. Lipid rafts in lymphocyte activation and migration. Mol. Membr. Biol. 23, 59–69 (2006).

Masedunskas, A., et al. Activated leukocyte cell adhesion molecule is a component of the endothelial junction involved in transendothelial monocyte migration. FEBS Lett. 580, 2637–2645 (2006).

Mitchell, D. M., Lovel, A. G. & Stenkamp, D. L. Dynamic changes in microglial and macrophage characteristics during degeneration and regeneration of the zebrafish retina. J. Neuroinflammation. 15, 163 (2018).

Mitra, S., et al. Vegf signaling between Muller glia and vascular endothelial cells is regulated by immune cells and stimulates retina regeneration. Proc. Natl. Acad. Sci. U. S. A. 119, e2211690119 (2022).

Nagashima, M., Barthel, L. K. & Raymond, P. A. A self-renewing division of zebrafish Muller glial cells generates neuronal progenitors that require N-cadherin to regenerate retinal neurons. Development 140, 4510–4521 (2013).

Nagashima, M. & Hitchcock, P. F. Inflammation regulates the multi-step process of retinal regeneration in zebrafish. Cells 10, 783 (2021).

Nummer, D., et al. Role of tumor endothelium in CD4+ CD25+ regulatory T cell infiltration of human pancreatic carcinoma. J. Natl. Cancer. Inst. 99, 1188–1199 (2007).

Ofori-Acquah, S. F., King, J., Voelkel, N., Schaphorst, K. L. & Stevens, T. Heterogeneity of barrier function in the lung reflects diversity in endothelial cell junctions. Microvasc. Res. 75, 391–402 (2008).

Ott, H., Diekmann, H., Stuermer, C. A. O. & Bastmeyer, M. 2001. Function of Neurolin (DM-GRASP/SC-1) in guidance of motor axons during zebrafish development. Developmental Biology 235, 86–97 (2001).

Rein, D. B., et al. Prevalence of Age-Related Macular Degeneration in the US in 2019. JAMA Ophthalmol. 140, 1202–1208 (2022).

Smith, J. R., Chipps, T. J., Ilias, H., Pan, Y. & Appukuttan, B. Expression and regulation of activated leukocyte cell adhesion molecule in human retinal vascular endothelial cells. Exp. Eye. Res. 104, 89–93 (2012).

Swart, G. W. Activated leukocyte cell adhesion molecule (CD166/ALCAM): developmental and mechanistic aspects of cell clustering and cell migration. Eur. J. Cell Biol. 81, 313–321 (2002).

Thummel, R., Bailey, T. J. & Hyde, D. R. In vivo electroporation of morpholinos into the adult zebrafish retina. J. Vis. Exp. 58, e3603 (2011).

Tudor, C., et al. Syntenin-1 and ezrin proteins link activated leukocyte cell adhesion molecule to the actin cytoskeleton. J. Biol. Chem. 289, 13445–13460 (2014).

Vihtelic, T. S. & Hyde, D. R. Light-induced rod and cone cell death and regeneration in the adult albino zebrafish (Danio rerio) retina. J. Neurobiol. 44, 289–307 (2000).

von Lersner, A., Droesen, L. & Zijlstra, A. Modulation of cell adhesion and migration through regulation of the immunoglobulin superfamily member ALCAM/CD166. Clin. Exp. Metastasis 36, 87–95 (2019).

Zhou, C., et al. Visual impairment and blindness caused by retinal diseases: A nationwide register-based study. J. Glob. Health 13, 04126 (2023).

