## Supplementary figure for "Alcama expressed in blood retina barrier and Muller glia is involved in zebrafish retina regeneration"

**Supplementary figures and table:**

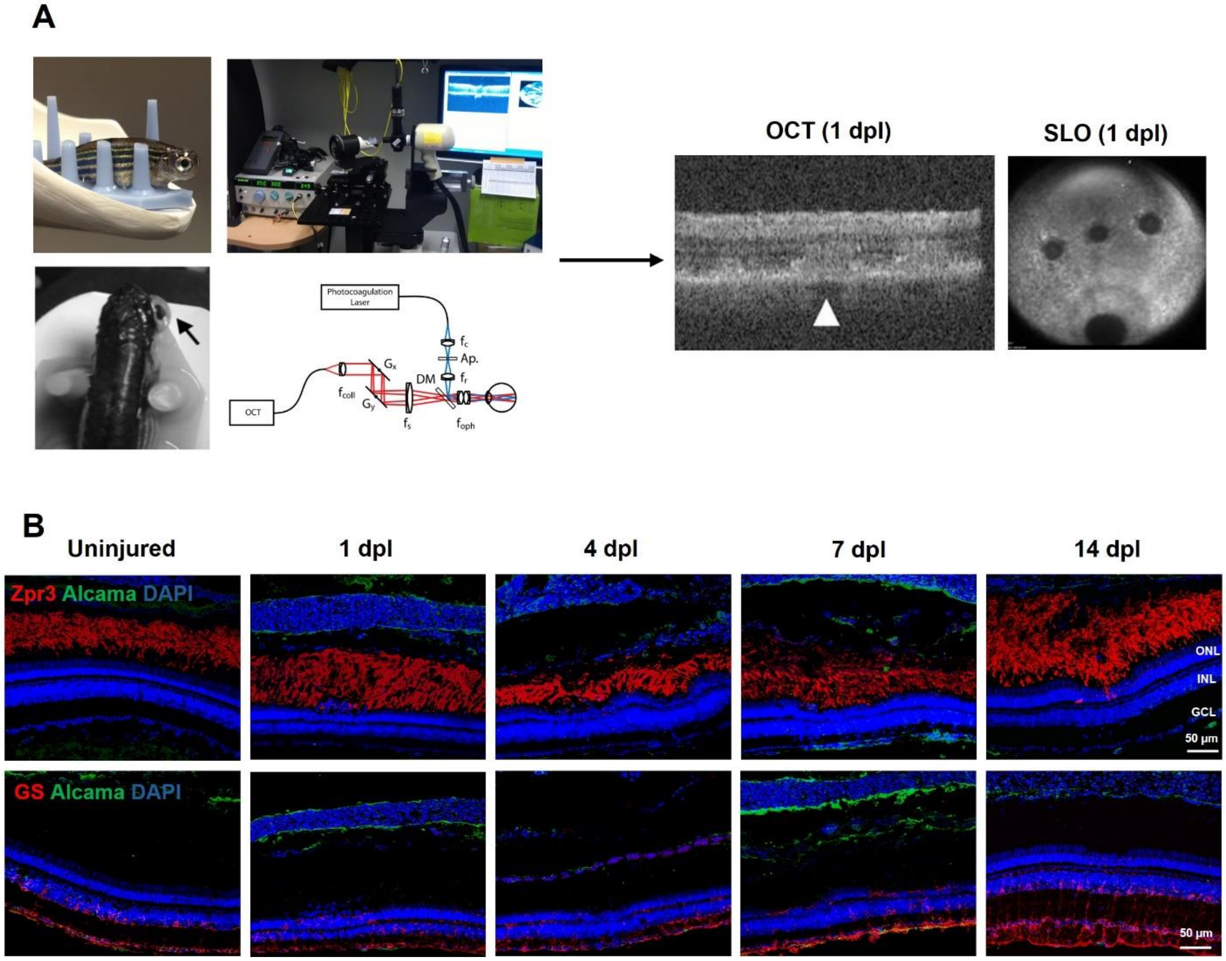


**Supplementary figure 1: (A)** Study design showing the instrument assembly for optical coherence tomography guided laser photocoagulation (Adapted from DiCicco, et al., 2014) and confirmation of lesion in the zebrafish retina using OCT and cSLO at 1 day post lesion (dpl). Arrowhead in OCT image points to a lesion spot where the photoreceptor layer has been ablated. SLO images showing three laser lesion spots superior to the optic nerve. **(B)** Representative IHC images showing Alcama (green) with Zpr3 (red) that stains the rod photoreceptors (top panel) and GS (red) which stains the glutamine synthetase present in Muller glial cells (bottom panel). Dapi (blue) indicates the nuclear staining. GCL-ganglion cell layer, INL-inner nuclear layer; ONL-outer nuclear layer.

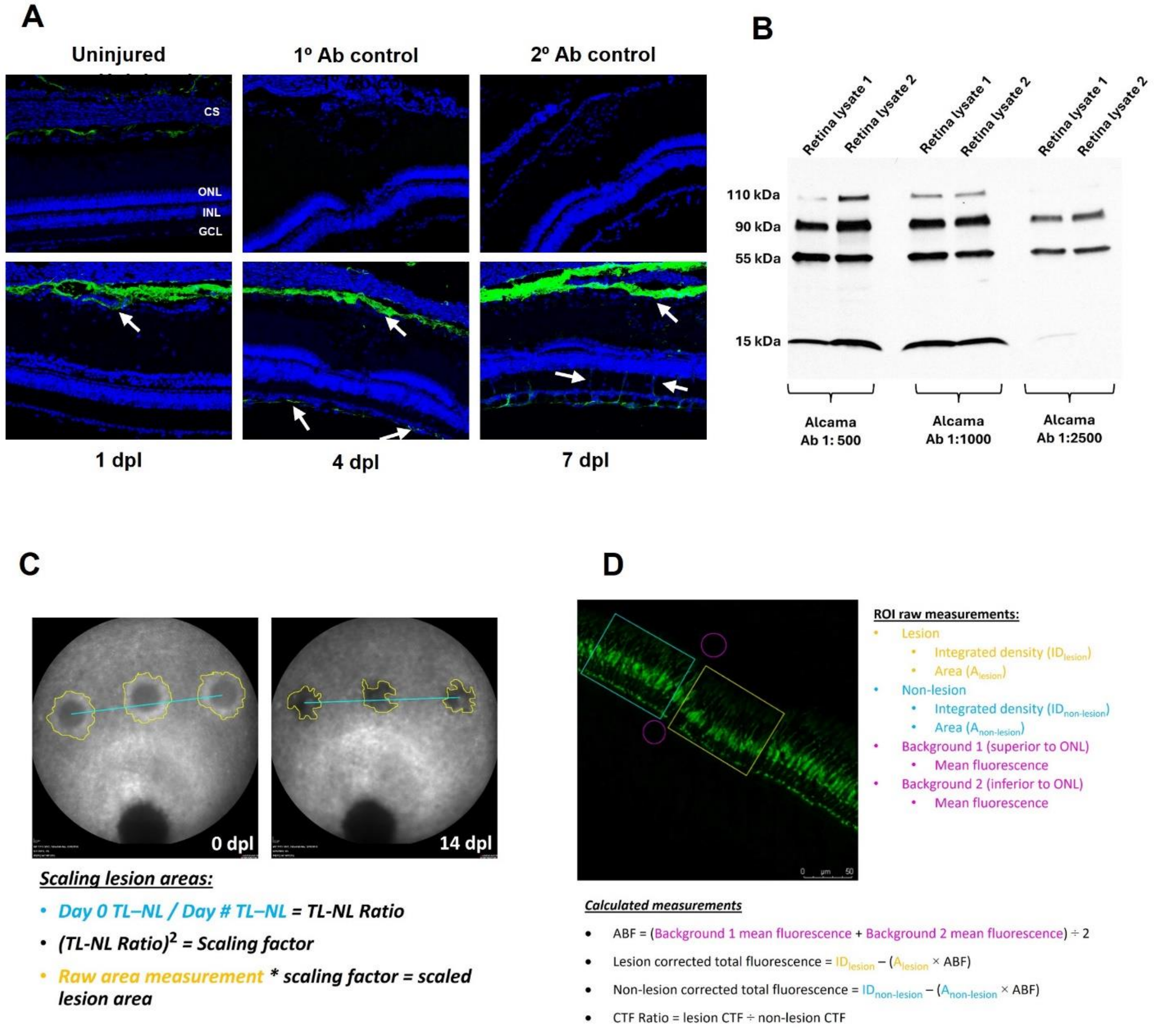


**Supplementary figure 2: (A, B)** Specificity of Alcama antibody was checked using Western blot and immunohistochemistry. **(C)** SLO imaging analysis quantification strategy. SLO images were used to track lesion areas in the same fish over time. Example images are shown for 0 dpl and 14 dpl to demonstrate the measurement of lesion areas and calculation of scaled lesion areas. Lesion traces (yellow) were made around the outermost edge of a lesion, discernible as different from the surrounding area. Temporal lesion to nasal lesion lengths (TL-NL; cyan) were measured and used to calculate the scaling factor to obtain more accurate scaled lesion areas. **(D)** Zpr1 (cone density) quantification strategy. Example image of section stained with Zpr1 antibody (green, labeling cone photoreceptor cells) is shown to demonstrate quantitative approach taken with each section. Lesioned (yellow) and non-lesioned (cyan) areas were drawn to obtain integrated density (ID) and area (A) measurements; background measurements also taken for each image (magenta). Calculations shown for average background fluorescence (ABF), corrected total fluorescence (CTF), and the ratio of CTF measurements in lesioned vs. non-lesioned areas to obtain the proportion of cones regenerated relative to the baseline cone density for each section/image.

**Table 1: List of antibodies used**

**Primary Antibodies**

| **Antibody Name** | **Catalog #** | **Source** | **Host** | **Dilution** |
|---|---|---|---|---|
| Alcama | GTX128399 | GeneTex | Rabbit | 1:100 |
| GFAP | ZDB-ATB-081002-46 | ZFIN | Mouse | 1:100 |
| Glutamine Synthetase | 66323-1-Ig | Proteintech | Mouse | 1:500 |
| Huc/D | A-21271 | Invitrogen | Mouse | 1:50 |
| PCNA | SC-56 | Santa Cruz Biotechnology | Mouse | 1:100 |
| ZPR-1 | ZDB-ATB-081002-43 | ZFIN | Mouse | 1:5 |
| ZPR-3 | ZDB-ATB-081002-45 | ZFIN | Mouse | 1:200 |
| 4C4 | N/A | generated in-house | Mouse | 1:500 |

**Secondary Antibodies**

| **Antibody Name** | **Catalog #** | **Source** | **Conjugate** | **Host** | **Dilution** |
|---|---|---|---|---|---|
| Anti-Mouse IgG | A-11004 | ThermoFisher Scientific | Alexa Fluor 568 | Goat | 1:200 |
| Anti-Rabbit IgG | A-11008 | ThermoFisher Scientific | Alexa Fluor 488 | Goat | 1:200 |
| Anti-Rabbit IgG | A-11011 | ThermoFisher Scientific | Alexa Fluor 568 | Goat | 1:200 |
